# Resting-state functional connectivity predictors of subjective visual Gestalt experience

**DOI:** 10.1101/2022.10.12.511888

**Authors:** Marilena Wilding, Anja Ischebeck, Natalia Zaretskaya

## Abstract

Subjective perceptual experience is influenced not only by bottom-up sensory information and experience-based top-down processes, but also by an individual’s current brain state. Specifically, a previous study found increased prestimulus insula and intraparietal sulcus (IPS) activity before participants perceived an illusory Gestalt (global) compared to the non-illusory (local) interpretation in a bistable stimulus. This study provided only a snapshot of the prestimulus brain state that favors an illusory interpretation. In the current study, we tested the hypothesis that the neural machinery that biases perception towards the illusory interpretation immediately before the stimulus onset, is also predictive of an individual’s general tendency to perceive it, which remains stable over time. We examined individual differences in task-free functional connectivity of insula and IPS and related it to differences in the individuals’ duration of the two interpretations. We found stronger connectivity of the IPS with areas of the default mode and visual networks to predict shorter local perceptual phases, i.e., a faster switch to an illusory percept, but no equivalent results for the insula. Our findings suggest a crucial role of an IPS interaction with nodes of key intrinsic networks in forming a perceptual tendency towards illusory Gestalt perception.

## 1. Introduction

Our whole awake life, the brain is engaged in interpreting the overwhelming amount of visual input. We constantly select, sort and group the incoming information to create a meaningful experience of the external world. These processes are usually automatic and are only called into question when an ambiguous or noisy cue has been interpreted in an illusory way rather than by its mere physical properties, e.g., when we misinterpret an object shadow as a threatening creature in the darkness or see contours of animals in cloud formations. These examples illustrate that the subjective interpretation of the stimuli is determined not only by the bottom-up sensory information that comes into the visual system, but also by our knowledge, goals and expectations, which shape the outcome of perception in a top-down fashion.

The neural mechanisms of these subjective, interpretative aspects of perception are often investigated using bistable (sometimes also called ‘ambiguous’) stimuli. These visual stimuli can be interpreted in two different ways based on the same sensory input. During continuous viewing, subjective perception alternates spontaneously between these mutually exclusive interpretations, a state which is known as *perceptual bistability* (Sterzer et al. 2009; Martínez and Parra 2018; Devia et al. 2022). A special type of bistable stimuli, termed ‘asymmetric’ bistable stimuli, is particularly relevant for studying the interpretative processes in perception. In contrast to typical bistable stimuli, which contain two equally complex perceptual interpretations, these stimuli yield one simpler interpretation, that is closer to the actual sensory input, and one more complex, illusory interpretation, that is derived from the simpler one (Lorenceau and Shiffrar 1992; Anstis and Kim 2011).

Previous research with bistable stimuli showed that subjective perceptual experience is influenced not only by the sensory input and top-down influences, but also by an individual’s current brain state (Kleinschmidt et al., 2012). For example, spontaneous brain activity in areas specialized for representing a certain sensory content increases before an ambiguous stimulus is perceived according to the area’s ‘preferred’ interpretation. Specifically, spontaneous activity in the human face-selective fusiform face area is higher if participants report perceiving a face in the ambiguous face-vase illusion (Hesselmann et al. 2008). In a recent fMRI study, we could demonstrate a similar effect using an asymmetric bistable stimulus with a *global* (illusory) and a *local* (non-illusory) interpretation (Anstis illusion, Anstis and Kim, 2011; Wilding et al., 2022).

We found increased prestimulus insula and intraparietal sulcus (IPS) activity before participants perceived the *global* compared to the *local* interpretation. These two key regions are repeatedly implicated in bistable perception paradigms (Sterzer and Kleinschmidt 2010; Zaretskaya et al. 2013; Grassi et al. 2018). Since this study used a paradigm with task-free prolonged fixation periods interleaved with brief stimulus presentations, the findings suggest that an upcoming percept may be determined (or at least biased) by signal fluctuations in intrinsic resting-state brain networks. Specifically, prestimulus insula activity seems to indicate the involvement of the salience network and the IPS activity the involvement of the dorsal attention network in favoring the illusory *global* percept.

Importantly, the study discussed above provided only a snapshot of the prestimulus brain *state* that leads to favoring the illusory interpretation of the visual input. However, the perception of a bistable stimulus fluctuates not only moment-to-moment within an individual. There are considerable inter-individual differences in the parameters of bistable perception across individuals, which remain relatively stable over time in conventional as well as asymmetric bistable paradigms (Kleinschmidt et al. 2012; Boeykens et al. 2019). Previous studies successfully linked inter-individual variability in the reversal rate of conventional bistable stimuli to various aspects of brain structure and function, such as cortical thickness (Kanai et al. 2010, 2011), microstructural gray matter tissue properties (Kanai et al. 2010), and resting-state connectivity (Baker et al. 2015; Mao et al. 2020). These findings suggest that an individual’s dynamic of bistable perception represents a *trait-like* property with corresponding signatures on the neural level.

In the current study, we tested whether the neural machinery that predicts the illusory *global* percept occurrence, is also predictive of a subject’s individual tendency to perceive the illusory component of the sensory input. We addressed this research question by examining individual differences in task-free resting-state functional connectivity of the superior insula and the anterior IPS regions, and linking them to the individual differences in the average duration of the *local* and *global* perceptual phases of an asymmetric bistable stimulus. To ensure that observed resting-state functional connectivity signatures are stimulus-specific and are not inherent to all bistable stimuli, we used three additional bistable stimuli with different sensory content and involving different perceptual mechanisms. We found that stronger connectivity of the IPS with the core default mode network (DMN) as well as with extrastriate visual areas predicted shorter *local* perceptual phases, i.e., a faster switch to an illusory, *global* percept. We did not find any resting-state signature of the *global* phase duration after the switch from local, and no equivalent results for the insula. Our findings show that several key nodes across different intrinsic networks may work together in shaping the self-generated illusory aspects of perceptual Gestalt experience. They also suggest that the overall tendency to perceive the illusory aspect of the stimulus is driven by two distinct mechanisms, one facilitating a switch to the *global* percept, and the other maintaining the *global* percept in consciousness, with the latter not being captured by the resting-state connectivity patterns.

## 2. Material and Methods

### 2.1 Participants

56 healthy volunteers (*m*= 24.0, *SD*= 2.83; 34 female) were included in this study. Four participants were excluded from the original sample of 60 participants due to technical difficulties during respiration recording or the inability to perceive both options of at least one bistable stimulus. All participants were right-handed and had no neurological, cardiovascular, or psychiatric diseases. They had normal or corrected to normal visual acuity and were not taking any medication regularly. Participants received a standardized instruction before they gave written informed consent. They received monetary compensation. Psychology students could alternatively choose compensation in the form of course credit for their participation. The study was approved by the local ethics committee of the University of Graz and was conducted according to the general principles of the Declaration of Helsinki.

### 2.2 Data acquisition

#### MRI session

Resting-state data were acquired on a 3T MRI system (MAGNETOM Vida, Siemens, Erlangen, Germany), with a 64-channel head coil. First, a structural T1-weighted image was acquired for each participant, using the following parameters: 176 slices, voxel size= 1 × 1 × 1 mm, TR= 1530 ms, TE= 3.88 ms, TI= 1200 ms, flip angle= 7 °, FOV= 256 mm. Subsequently, resting-state functional data were acquired in an 8-minute session, in which participants were instructed to fixate a central red dot on the screen and to avoid directed thoughts. Physiological signals were recorded during the whole resting-state period, using a photoplethysmograph (cardiac signal) and a respiratory belt, both with a sampling rate of 400 Hz. This was done to avoid contamination of functional data with physiological noise, which is especially severe in resting state functional connectivity (RSFC) studies, as it introduces spurious correlations (Deckers et al. 2006; Chang and Glover 2009; Birn 2012; Murphy et al. 2013). The functional data were acquired using a T2*-weighted 2D echo-planar imaging sequence with the following parameters: 46 interleaved slices, no slice gap, voxel size= 3 × 3 × 3 mm, repetition time (TR) = 3220 ms, echo time (TE) = 32 ms, flip angle= 82 °, FOV = 228 mm, 155 volumes. After collecting resting-state data, which is the focus of this study, subsequent functional, diffusion and structural scans were acquired that are not part of the current research question.

#### Behavioral session

To avoid the impact of experience with visual illusions on the resting-state scan, behavioral data was collected on a separate day after the fMRI session, with two sessions being on average 11.54 days apart (SD = 15.87) apart. Bistable stimuli were presented on a gamma-corrected 24-inch Asus VG248QE LCD gaming monitor (ASUSTeK Computer Inc., Taipwei, Taiwan) with a refresh rate of 60 Hz, a resolution of 1920 × 1080 mm (44.44 × 25.91° v.a.) and a maximum luminance of 90.4 cd/m^2^, using Psychtoolbox-3 (Kleiner et al. 2007) and MATLAB 2017b (MathWorks, Inc., Natick, MA) running under Linux Ubuntu 18.04 LTS. During the whole experiment, participants rested their head on a chin rest, which was positioned approximately 65 cm from the monitor and were asked to fixate the red dot in the center of the screen (dot size: 0.27° v.a.). Each stimulus was presented four times for 120 s, interleaved with a 20 s baseline period. The order of the stimuli was pseudorandomized and counterbalanced across participants. During stimulus presentation, participants were instructed to continuously indicate which of the two interpretations they perceive by pressing either the left or the right button on a high-precision mechanical gaming keyboard (APEX M800, SteelSeries, Denmark) for as long as they experienced one interpretation. Whenever their percept switched, they changed to the other button, letting go of both buttons only briefly during switching (or when they were unsure of their percept). Before each run, participants familiarized themselves with the two possible interpretations of each bistable stimulus in an exercise run lasting 30-60 s. This resulted in a total time of around 9 minutes per run/illusion. Over all illusions and participants, the average total measuring time was 45.22 min. (*SD*= 4.10).

#### Bistable stimuli

The Anstis global-local illusion (Figure 1) consists of eight dots (dot width: 0.41° v.a.) organized in pairs, with each pair rotating around one of the four imaginary axes with a radius of 0.62° v.a. in a 2D plain at 0.5 revolutions per second. When viewed continuously, the stimulus can be perceived either as four rotating dot pairs (*local* non-illusory interpretation) or as two moving illusory squares (*global* illusory interpretation) with a side length of 5.89° v.a. that are formed by long-range grouping of the individual dots (Anstis and Kim 2011). It is well-known that parameters of different bistable stimuli correlate with each other, with the degree of correlation depending on the similarity of the involved mechanisms (Carter and Pettigrew 2003; Gallagher and Arnold 2014; Cao et al. 2018). Therefore, to determine which of the effects are specific to the illusory long-range grouping in the Anstis illusion, and which are shared with other types of bistable perception, we used three additional bistable stimuli which are also illustrated in Figure 1. The Rubin face-vase illusion (side length: 6.9° v.a.) can appear either as two faces, which are turned to each other or one vase in the middle (Rubin 1915). In the Coffer illusion (side length: 6.9° v.a.), one can either see several rows of squares lined up next to each other or have the impression of several circles suddenly appearing in front of the squares (Norcia 2006). Finally, in the ambiguous structure-from-motion (SFM) illusion (350 black dots with dot size 0.16° v.a., randomly placed within an aperture of 3.15° by 6.14° and moving at a speed of 0.56°v.a./s in opposite directions) one’s perception changes between leftward and rightward rotation of a 3D cylinder (Wallach and O’Connell 1953). The stimulus set was put together to engage either the dorsal or the ventral processing streams (moving and static, respectively) and to contain either two equally complex interpretations (SFM, Rubin) or one illusory and one more simple interpretation (Anstis, Coffer).

**Figure 1.**
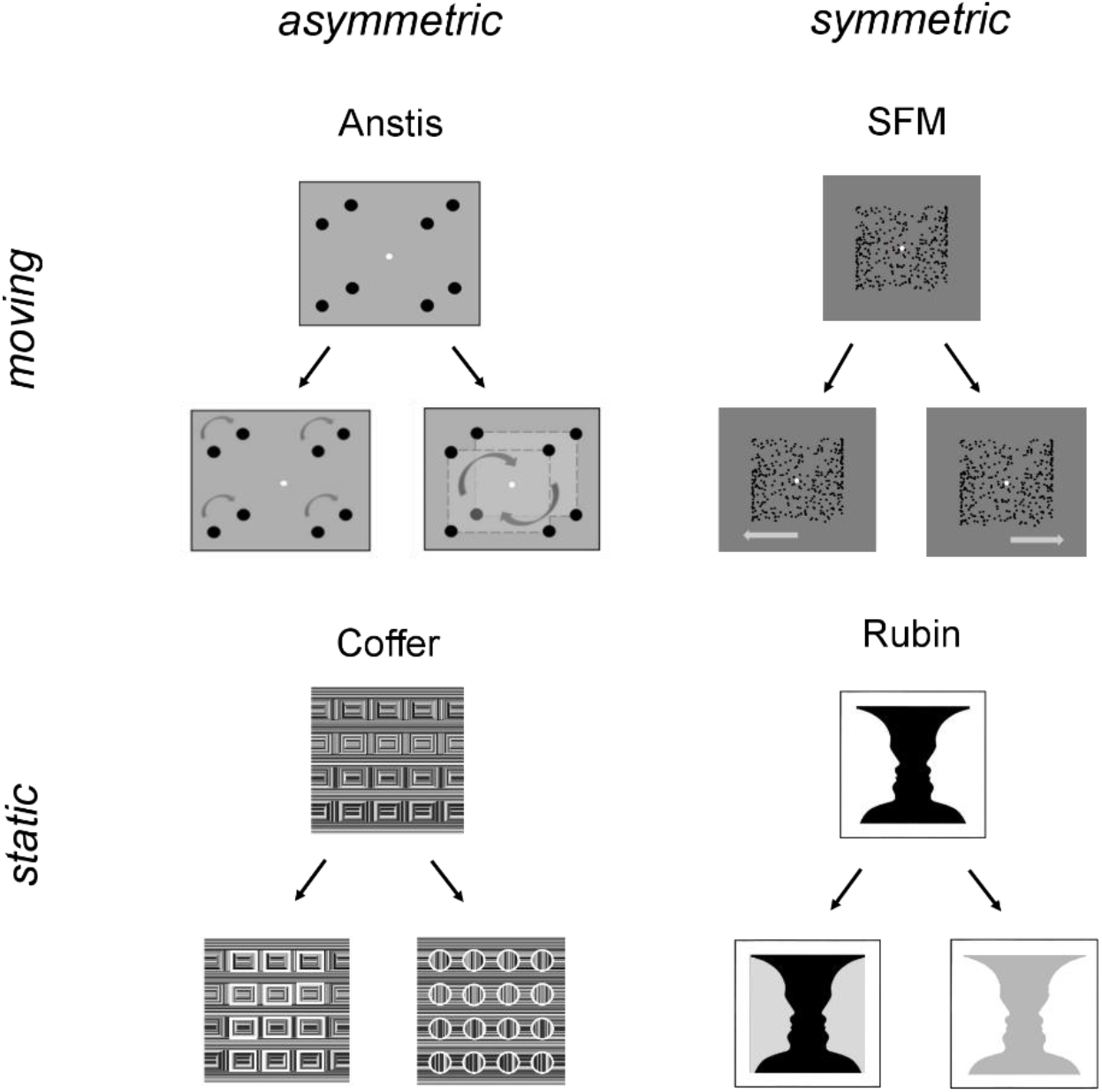
Illustration of the two alternative percepts for each of the four bistable illusions, separated in static and moving as well as asymmetric and symmetric stimuli.

### 2.3 Data analysis

#### Behavioral data analysis

Analysis of behavioral data was performed using custom scripts written in MATLAB version R2019b (MathWorks, Inc., Natick, MA). Subjects’ button presses during illusion perception were used to infer the identity and duration of each perceptual state. Individual’s percept durations in bistable perception as well as their means in the population are known to follow a left-skewed distribution (Levelt 1967; Brascamp et al. 2005; Moreno-Bote et al. 2007; Mao et al. 2020). Therefore, to avoid biasing the correlation/regression analysis by extreme values, we calculated the geometric mean dominance duration, defined as the n^th^ root of the product of n numbers, instead of an arithmetic mean. We did this separately for each percept of each illusion. To investigate the relation between and within illusions/percepts, we performed Spearman’s rank correlation analysis.

#### Preprocessing of (f)MRI data

Structural and functional data were preprocessed using fMRIPrep version 20.2.3 (Esteban et al. 2019), based on Nipype 1.6.1 (Gorgolewski et al. 2011). Parts of the following descriptions were taken from the fMRIPrep boilerplate text and modified.

For the anatomical preprocessing, the T1-weighted image was corrected for intensity non-uniformity and then used as a reference scan for the subsequent steps. The T1w-reference was skull-stripped and then brain tissue was segmented in white matter (WM), gray matter (GM) and cerebrospinal fluid (CSF). Next, brain surfaces were reconstructed using FreeSurfer’s recon-all pipeline (FreeSurfer 6.0.1; Dale et al., 1999). For the functional data, a skull-stripped reference volume was generated. Susceptibility distortion was estimated from the corresponding images with reversed phase-encoding and a corrected EPI image was computed. This reference was co-registered with the T1w image using boundary-based registration with 6 degrees of freedom (Greve and Fischl 2009). Head motion parameters (transformation matrices, and six corresponding rotation and translation parameters) were estimated before any spatiotemporal filtering was applied using *mcflirt* (FSL 5.0.9). Next, transformations related to head motion correction, slice-time correction (AFNI), susceptibility distortion correction and remapping onto the fsaverage standard space were concatenated and applied in one interpolation step. The output of fmriprep was smoothed on the surface using a 2D Gaussian kernel with the full-width at half-maximum (FWHM) of 5 mm.

#### Functional connectivity analysis

In order to determine the resting-state correlates of bistable perception, we computed seed-based resting-state functional connectivity (RSFC) with superior insula and anterior IPS as seeds, as we found these two regions to be relevant in our previous study (Wilding et al. 2022). The anterior IPS was defined using the probabilistic atlas of visual topography (IPS5; Wang et al., 2015). The superior insula ROI was taken from the Destrieux parcellation atlas provided by FreeSurfer (Fischl et al. 2004). Analysis was performed using FreeSurfer and FS-FAST version 7.1.0.

For the first-level single-subject analysis, the average time course of each of the two seed regions, anterior IPS and the superior insula, was extracted for each hemisphere and then averaged over both hemispheres. The time course of the respective seed was included in the GLM as regressor of interest. In addition, we used the physiological signals recorded during the resting state scan to account for physiological noise. Pulse and respiration signals were processed using the MATLAB-based PhysIO toolbox (Kasper et al. 2017) as described in our previous work (Wilding et al., 2022). In brief, the physiological data were aligned to the fMRI time series, preprocessed using a peak detection algorithm and subsequently RETROICOR phase expansion (Glover et al. 2000) was applied. The 18 resulting regressors were then downsampled to a reference slice of each volume and added to the GLM as nuisance regressors. To account for head-motion induced artifacts, six motion regressors were added to the GLM.

Individual beta estimates for the seed time course regressor, which represent the functional connectivity of the seed with each cortical surface location, were taken to the second-level group analysis. The group analysis consisted of the whole-brain regression with mean dominance durations (geometric mean) of each percept as predictor (X) and connectivity maps as data (Y). Group results were corrected using the Monte Carlo Simulation method with a cluster-forming threshold of p<0.01 and a cluster significance level of p< .05 (two-sided), with an additional Bonferroni correction for two spaces (left and right hemisphere surface) of the cluster-level p-value. Attribution of significant clusters to the resting-state networks was performed using the Schafer parcellation with 400 parcels (Schaefer et al. 2018).

Our previous study showed that a joint activity increase in the IPS and the insula predicts the subsequent global perception of an individual (Wilding et al. 2022), suggesting perception-related covariation of activity in the two areas. Therefore, in addition to performing a seed-based functional connectivity analysis for each seed separately, we performed the unshared functional connectivity analysis by adding time courses of both seeds into the same first-level GLM. This allowed us account for resting-state functional connectivity that is common to the two seeds and examine their unique connectivity targets.

## 3. Results

### 3.1 Behavioral results

Since the dominance durations of different bistable stimuli typically correlate with each other (Carter and Pettigrew 2003; Gallagher and Arnold 2014; Cao et al. 2018), prior to the main analysis we first analyzed the correlation structure of dominance durations in our dataset. This information is important because strongly correlated dominance durations may manifest themselves as similar connectivity maps in our second-level group analysis (see below).

For all but the Anstis illusion, strongest correlations were observed for dominance durations between percepts within the same illusion (Table 1). Interestingly, we did not observe a significant association between the durations of the illusory (*global*) and non-illusory (*local*) interpretation of the Anstis illusion (*r*= 0.26, *p*= 0.06), suggesting that the two perceptual interpretations in this stimulus are governed by independent perceptual and neural mechanisms (Figure 2). Consistent with previous findings, durations of percepts belonging to different illusions showed weak to moderate correlations (Cao et al. 2018).

**Table 1.** Correlations of the geomean durations between all interpretations of all four illusions.

|  | *local* | *global* | *face* | *vase* | *square* | *circle* | *left* |
| --- | --- | --- | --- | --- | --- | --- | --- |
| *local* | - | - | - | - | - | - | - |
| *global* | ***r= 0.26*** | - | - | - | - | - | - |
| *face* | r= 0.31* | r= 0.30* | - | - | - | - | - |
| *vase* | r= 0.40** | r= 0.34* | ***r= 0.68\*\*\**** | - | - | - | - |
| *square* | r= 0.36** | r= 0.18 | r= 0.54*** | r= 0.46*** | - | - | - |
| *circle* | r= 0.39** | r= 0.41** | r= 0.57*** | r= 0.62*** | ***r= 0.73\*\*\**** | - | - |
| *left* | r= 0.13 | r= 0.33* | r= 0.26 | r= 0.26 | r= 0.21 | r= 0.41** | - |
| *right* | r= 0.20 | r= 0.21 | r= 0.28* | r= 0.47*** | r= 0.34* | r= 0.42** | ***r= 0.65\*\*\**** |

Note. \* p < .05, \*\* p < .01, \*\*\* p < .001. Intra-stimulus correlations are shown in ***italic bold font***.

**Figure 2.**
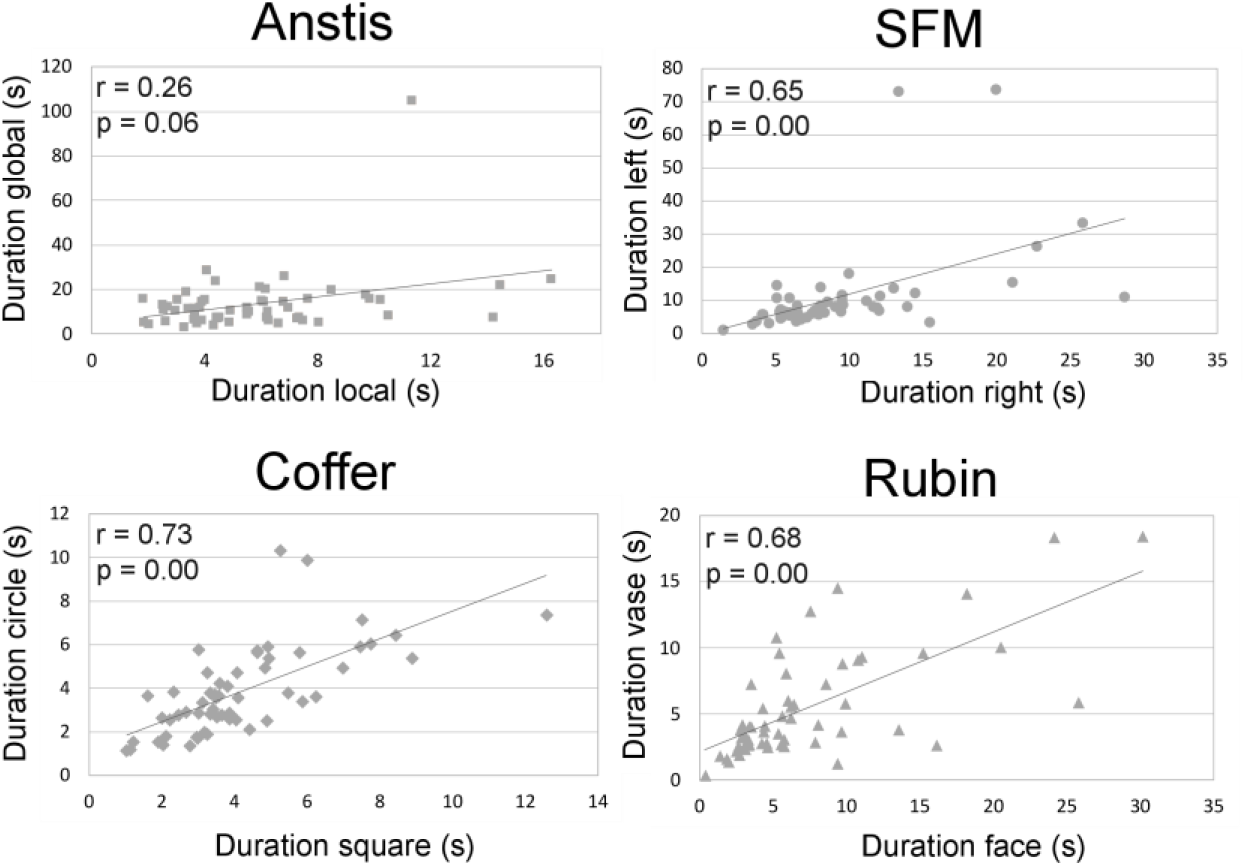
Correlations between geomean durations of the two interpretations for each bistable illusion. SFM, Coffer and Rubin’s illusions show strong correlations between percepts. The durations for the *local* and *global* interpretation in the Anstis illusion display a much weaker association that accordingly does not reach statistical significance.

### 3.2 Functional connectivity results

#### 3.2.1 Salience, Dorsal Attention and Frontoparietal network

We first confirmed that our seed regions identified based on findings of our previous study (Wilding et al. 2022) engaged hypothesized intrinsic resting-state networks, as illustrated by the group connectivity results of each seed in Figure 3 and Table 2. As expected, the insula seed exhibited functional connectivity primarily with the salience network, and the anterior IPS seed with the frontoparietal network comprised of the dorsal attention and frontoparietal control networks (DAN and FPCN) as defined by the 400 parcel Schaefer parcellation (Schaefer et al. 2018).

**Figure 3.**
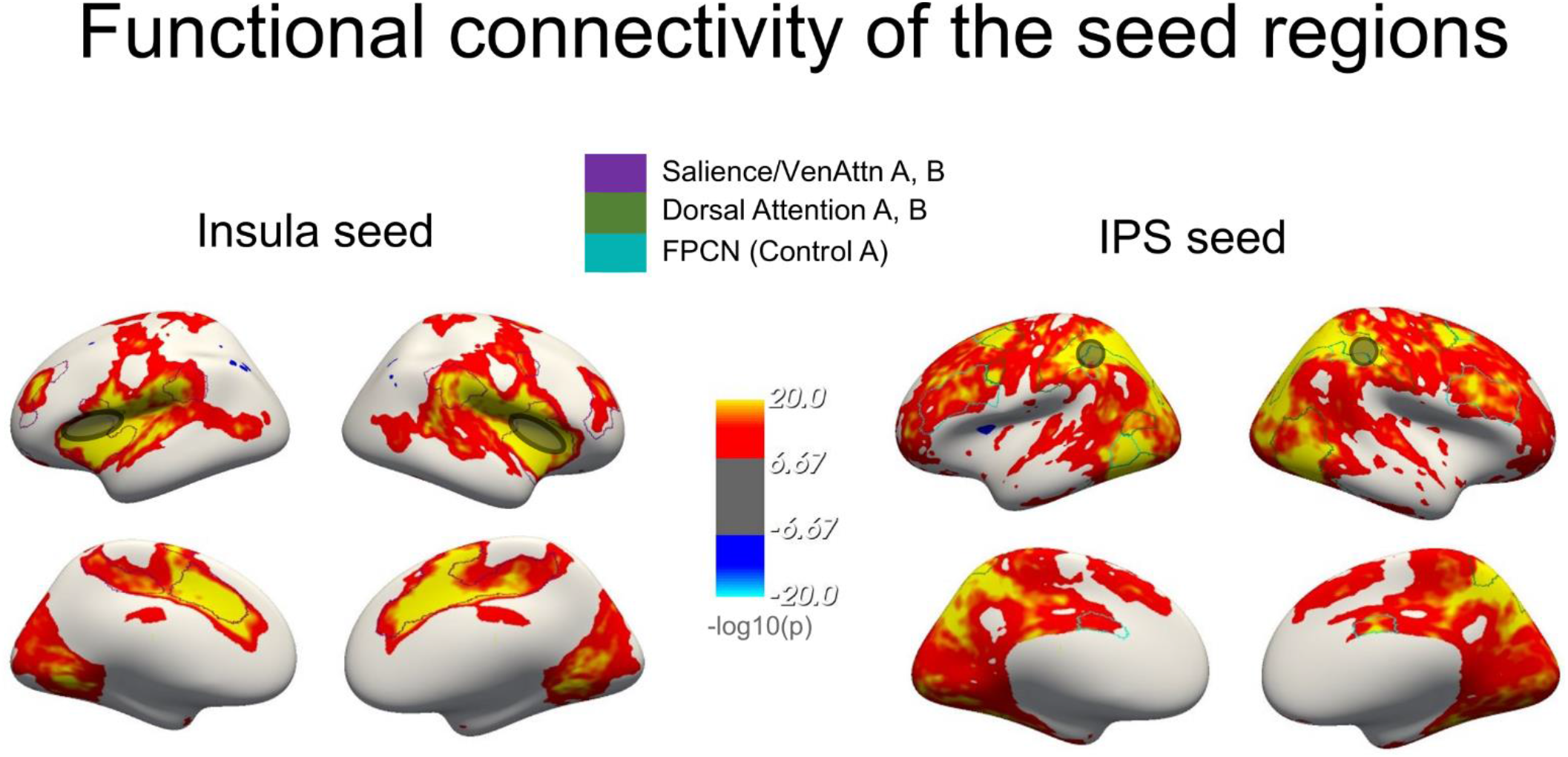
Resting-state networks that exhibit functional connectivity with the respective seed regions. The superior insula displayed significant FC with the salience network (violet) whereas the IPS showed strong connections with the dorsal attention (green) and frontoparietal control (turquoise) network. Seed regions are shown as an outlined ellipse/circle. Connectivity maps are shown at a cluster-forming threshold of p< 10^−8^ and a cluster-level p< 0.05 (corrected).

**Table 2.** Quantification of overlap (%) between functional connectivity of the seed regions with the salience, the dorsal attention and the control network in each hemisphere.

| *Seed* | *Hemi* | *SN* | *DAN* | *FPCN* |
| --- | --- | --- | --- | --- |
| Insula | left | **90.42** | 21.76 | 19.41 |
|  | right | **94.61** | 26.12 | 27.14 |
| IPS | left | 42.25 | **96.18** | **95.45** |
|  | right | 46.71 | **97.32** | **97.03** |

*Note.* SN = Salience network, DAN = Dorsal attention network, FPCN = Frontoparietal control network. Maximum overlap of each seed's connectivity and each network is shown in **bold font**.

#### 3.2.2 Insula seed

To address our research question, we first investigated whether there is a relationship between dominance durations of the *global* and *local* percepts in the Anstis illusion and the functional connectivity of the superior insula. The results of this analysis are shown in Figure 4A and Table 3. We found an association between shorter *local* percept durations in the Anstis illusion and insula’s connectivity strength with the left inferior frontal gyrus (IFG) as well as temporo-parietal regions.

**Figure 4.**
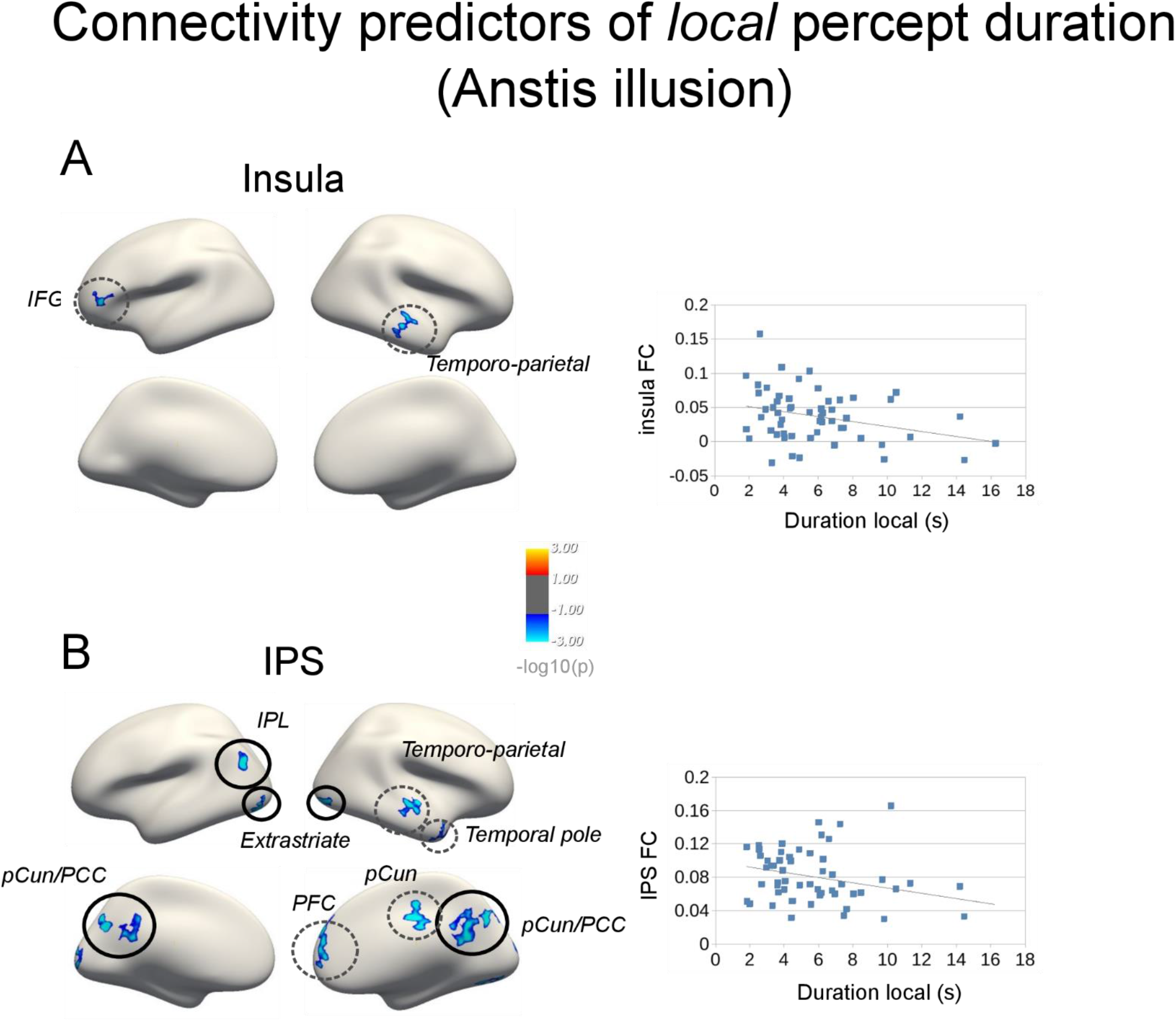
Resting-state functional connectivity predictors of percept duration with A) dorsal insula and B) anterior IPS as seed regions. Scatterplots show the direction of the relation between percept durations and FC of each seed region with the respective clusters (average FC over clusters); p< .01, corrected. Effects that are unique to a specific seed (i.e., are significant in unshared functional connectivity analysis, see below) are framed by black circles.

**Table 3.**
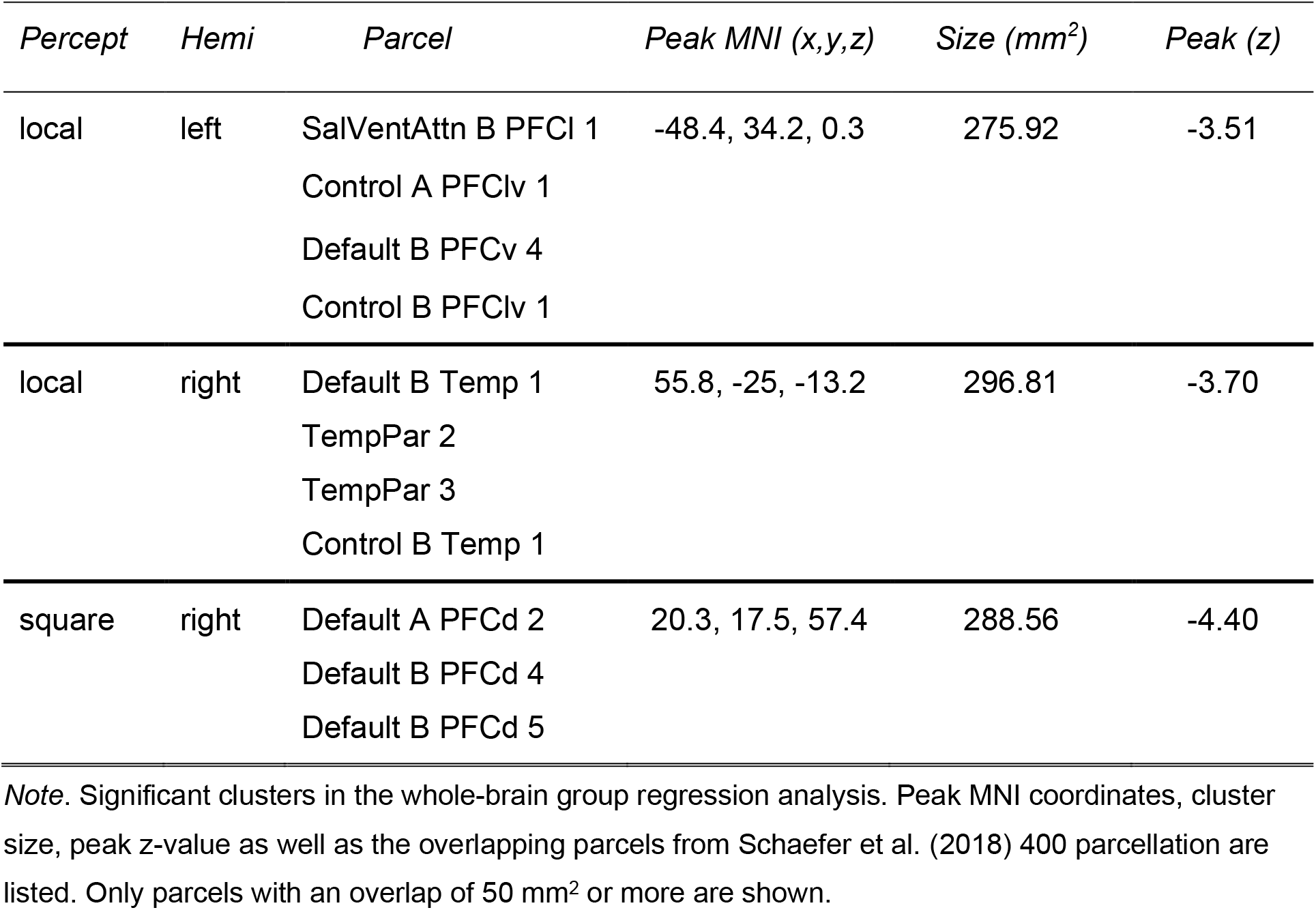
Functional connectivity predictors of individual’s percept duration (insula seed)

We then tested whether similar results could be found for the percept durations of other illusions. This analysis revealed a single association between shorter *square* percept durations of the Coffer illusion and insula’s connectivity strength with a cluster in superior frontal region (see Figure S1). There were no significant effects for other illusions.

#### 3.2.3 IPS seed

We further tested whether there is a relationship between percept durations of the Anstis illusion and the functional connectivity of the anterior IPS. We found a negative relationship between the *local* percept durations and the IPS connectivity strength with areas of the ‘core’ default mode network (Andrews-Hanna et al. 2014), namely within the right prefrontal cortex (PFC), the left inferior parietal lobule (IPL), as well as bilateral precuneus/posterior cingulate cortex (PCC) areas. Additionally, we observed significant negative associations with clusters in the right temporal lobe and the extrastriate cortex (see Figure 4B and Table 4).

**Table 4.** Functional connectivity predictors of individual’s percept duration (IPS seed)

| *Percept* | *Hemi* | *Parcel* | *Peak MNI (x.y.z)* | *Size (mm<sup>2</sup>)* | *Peak (z)* |
| --- | --- | --- | --- | --- | --- |
| local | left | Default A IPL 1 | -43.7, -68.6, 25.9 | 276.80 | -4.41 |
|  |  | Default A IPL 2 |  |  |  |
|  |  | Default C IPL 1 |  |  |  |
| local | left | VisCent ExStr 6 | -38.4, -86.0, -14.3 | 556.07 | -4.69 |
|  |  | VisCent ExStr 3 |  |  |  |
|  |  | VisCent ExStr 4 |  |  |  |
|  |  | DorsAttnA TempOcc 4 |  |  |  |
| local | left | VisCent ExStr 10 | -13.7, -93.6, 2.8 | 474 | -3.41 |
|  |  | VisCent Striate 1 |  |  |  |
|  |  | VisPeri StriCal 1 |  |  |  |
| local | left | ContC pCun 1 | -15.6, -65.2, 31.2 | 263.79 | -5.34 |
|  |  | ContC pCun 2 |  |  |  |
| local | left | Default A pCunPCC 1 | -4.4, -49.0, 21.6 | 378.11 | -3.25 |
|  |  | Default A pCunPCC 2 |  |  |  |
|  |  | Default A pCunPCC 3 |  |  |  |
| local | right | TempPar 2 | 62.3, -10.5, -3.0 | 356.60 | -5.10 |
|  |  | TempPar 3 |  |  |  |
|  |  | TempPar 4 |  |  |  |
| local | right | Default B AntTemp 1 | 36.0, 15.7, -35.1 | 380.62 | -3.62 |
|  |  | TempPar 1 |  |  |  |
|  |  | LimbicA TempPole 5 |  |  |  |
|  |  | LimbicA TempPole 3 |  |  |  |
| local | right | VisCent ExStr 5 | 43.1, -82.4, -8.4 | 242.99 | -4.28 |
|  |  | VisCent ExStr 6 |  |  |  |
| local | right | VisCent ExStr 10 | 22.1, -76.6, 43.4 | 398.04 | -4.84 |
| local | right | VisCent ExStr 3 | 24.6, -72.5, -5.8 | 307.61 | -4.08 |
| local | right | Default A pCunPCC 1<br>Default A pCunPCC 2<br>Default A pCunPCC 5<br>Default C Rsp 2<br>ContC pCun 2<br>DorsAttn A SPL 3<br>VisPeri ExStrSup 4 | 6.8, -59.3, 18.3 | 648.53 | -3.57 |
| local | right | Default A pCunPCC 4<br>SalVentAttnA ParMed 1<br>ConC Cingp 2 | 3.9, -15.6, 34.8 | 313.20 | -5.71 |
| local | right | Default B PFCd 1<br>Default A PFCm 4<br>Default A PFCm 3 | 7.1, 53.5, 30 | 461.57 | -3.79 |
| square | right | Default B PFCd 5<br>Default A PFCd 2<br>Default B PFCd 4 | 20.6, 17.0, 57.3 | 246.21 | -4.36 |

*Note.* Significant clusters in the whole-brain group regression analysis. Peak MNI coordinates, cluster size, peak z-value as well as the overlapping parcels from Schaefer et al. (2018) 400 parcellation are listed. Only parcels with an overlap of 50 mm<sup>2</sup> or more are shown.

Similar to insula’s connectivity, we tested whether similar results could be found for the percept durations of other illusions. We found a cluster in the superior frontal cortex associated with duration of the *square* percept in the Coffer illusion (Figure S1). There were no significant effects for other illusions.

The above results suggested that both insula and IPS connectivity strength inversely predicts the duration of the *local* percept, such that the stronger the connectivity, the shorter the *local* percept phase of an individual. They also suggest that the connectivity targets of the two areas that are predictive of the *local* percept duration differ. While the insula’s connectivity targets are located mainly in the IFG and a temporal cluster, the IPS connectivity targets primarily involve the extrastriate visual cortex, as well as the frontal, parietal, precuneus and the lateral temporal nodes within the DMN.

To test whether the above pattern of results is free of shared connectivity between seed regions, we conducted an additional regression analysis with ‘unshared’ functional connectivity of the insula and the IPS by adding the time courses of both seed regions into the same first-level GLM. This analysis, presented in Supplementary Figure S2, showed that the main connectivity clusters precuneus/PCC, IPL and the extrastriate cortex for the IPS remain stable, even after considering unique connectivity contribution of each seed region at every cortical location. For the insula, however, none of the original clusters remained significant, implying that most of the insula effects are due to its connectivity with the IPS.

## 4. Discussion

### Summary

In this study, we investigated the top-down interpretative processes involved in visual perception by linking resting-state connectivity of large-scale brain networks and an individual’s tendency to perceive the more complex, illusory interpretation of a bistable stimulus. We confirmed that especially the IPS, which is involved in biasing the perception towards the *global* illusory interpretation in task-based fMRI, is also relevant for determining the individual’s general propensity for the *global* perception in a *trait*-like manner. We found that interactions between the IPS and the core DMN, as well as the extrastriate cortex, predict how soon an individual abandons the *local* non-illusory interpretation and switches to the illusory alternative.

### Insula and IPS contributions to shaping illusory perception

In our previous study, we used a slow event-related paradigm with long inter-stimulus intervals and brief stimulus presentations to show that the *global* stimulus interpretation is preceded by increased activity within the superior insula and the IPS (Wilding et al. 2022). Our current findings extend these results by demonstrating that the IPS shapes the way participants perceive the Anstis stimulus not only in a moment-to-moment manner, but also as an individual *trait*. In other words, individual connectivity strength of this area with several nodes of the default mode and the visual network predicts the individual tendency for switching to an illusory percept. Importantly, these IPS connectivity targets remained predictive of the global percept duration even when insula connectivity with these areas was accounted for.

Our insula connectivity findings were less conclusive. Although insula connectivity with several areas was predictive of the local percept duration in our main analysis, these effects disappeared after accounting for IPS connectivity with the same areas. This suggests that insula effects could partially be explained by the covariation of insula resting-state activity with that of the IPS. It therefore seems that the IPS, and not the insula, is the primary driver of the resting-state connectivity predictors of the illusory Gestalt perception. This finding appears to be in contrast with our previous study that investigated immediate prestimulus predictors of illusion perception (Wilding et al. 2022). In that study, it was the insula rather than the IPS that showed the strongest and most consistent prestimulus effects. Taking the findings of both studies together, we conclude that the insula may be more involved in shaping subjective illusory perception in the current moment, while the IPS may drive the trait-like aspects of illusory perception that remain stable over time.

Our IPS results fit into a series of studies linking resting-state functional connectivity properties with different individual characteristics, such as working memory or executive control (Hampson et al. 2006; Seeley et al. 2007; Reineberg et al. 2015). Resting-state functional connectivity correlates of bistable perception dynamics have also been reported, with resting-state fluctuations and connectivity being predictive of switching speed with conventional bistable stimuli (Baker et al. 2015; Mao et al. 2020). In the current study we demonstrate that resting-state functional connectivity contains more than just the signature of perceptual stability or switching speed. Using a special type of bistable stimulus with an illusory and a non-illusory interpretation we were able to distill the specific connectivity signatures of self-generated, illusory aspects of perception, and specifically of the readiness of an individual to switch to an illusory interpretation of the sensory input.

### IPS connectivity targets

The anterior IPS is well-known to play a major role in bistable perception and in representing illusory aspects of perceptual experience (Britz et al. 2009; Kanai et al. 2010; Knapen et al. 2011; Zaretskaya et al. 2013; Grassi et al. 2018). An intriguing question is why specifically the interaction of the IPS with parts of the DMN, mainly the precuneus/PCC and the IPL, predicts how soon participants switch to the *global* percept. The DMN is typically thought to be responsible for self-generated and internally oriented cognition, being disengaged during any kind of sensory processing (Andrews-Hanna et al. 2014). However, multiple studies demonstrated that the DMN is active not only during rest, but also during sensory tasks requiring a disengagement from the current sensory input and/or a reliance on prior experience, such as delayed match to sample task or contextual priming (Smallwood et al. 2013; Konishi et al. 2015; Yeshurun et al. 2017; González-García et al. 2018; Murphy et al. 2018; Sormaz et al. 2018). The type of processing involved in such tasks has been termed ‘perceptual decoupling’ (Schooler et al. 2011). The results of this processing are subsequently integrated with the current sensory input.

Yeshurun and colleagues (2017) were able to demonstrate this notion of the DMN’s role in integrating extrinsic sensory information with prior intrinsic models. They provided subjects with diverging contextual information before presenting them with a story, which could consequently be interpreted in two different ways. Interestingly, neural activity within DMN regions was more similar within subjects sharing a certain interpretation of the story compared to subjects who interpreted the story in a different way. These findings suggest comparable neural patterns within the DMN in subjects sharing an interpretation of external events, which was guided by prior contextual information (Yeshurun et al. 2017). In light of these findings, we suggest that a faster switch towards *global* illusory perception in our study may be mediated by a similar DMN-based perceptual decoupling mechanism. Increased functional connectivity between the IPS and the DMN allows the brain to disengage from the current sensory input that is reflected in the *local* stimulus interpretation, and to produce a more complex internally generated *global* illusory percept.

Our finding that the DMN is involved in shaping subjective visual experience is thus consistent with the emerging view of the DMN being situated at the top of the processing hierarchy (Margulies et al. 2016). The DMN is thought to represent the most abstract and complex mental content, and is considered to be the source of experience and knowledge-based priors that are integrated with the current perceptual input (Smallwood et al. 2021; Yeshurun et al. 2021). It can therefore provide abstract information about the illusory percept to the visual areas and the IPS. Interestingly, similar interactions between the default-mode network and other intrinsic-networks are thought to underlie hallucinatory experiences (Alderson-Day et al. 2016), confirming the proposed commonalities between illusions and hallucinations (Pearson and Westbrook 2015).

Interestingly, apart from the DMN areas, we also found IPS connectivity with visual clusters of the extrastriate cortex to be predictive of how soon the *local* percept will be abandoned. These visual clusters are located outside of the primary visual cortex and are not a part of any topographic visual area (Wang et al. 2015). The location of these clusters is similar to the location of the lateral occipital complex (LOC), which is known for its role in processing real and illusory objects (Grill-Spector et al. 2001; Martinez et al. 2007; Cichy et al. 2011). Our results imply that connectivity of the IPS with the DMN on the one hand, and with the LOC on the other hand, together favor a faster switch to a more complex, grouped stimulus interpretation. While the IPS-DMN connectivity may represent the self-generated, internal aspect of the *global* percept, the IPS-LOC connectivity may represent its perceptual object-like qualities.

### No predictors for global percept duration

It is somewhat surprising that our analysis revealed primarily the resting-state correlates of the *local* percept duration but not the *global* one. Unlike most bistable stimuli that show a strong correlation between the durations of the two percepts across individuals (see Table 1 and Figure 2), the correlation between *global* and *local* percept duration for the Anstis illusion was unexpectedly weak. In a typical bistable paradigm, the two possible interpretations are thought to be governed by the same competitive mechanism of mutual inhibition and adaptation (Moreno-Bote et al. 2007, 2010; Shpiro et al. 2007; Alais 2012). The very weak (and not significant) correlation between percepts of the Anstis stimulus indicates that the mechanisms that determine the average duration for each of them may be different. It is possible that, while the duration of the *local* percept is determined by factors like adaptation and evidence accumulation, the ability of an individual to maintain the *global* percept in consciousness may be related to other factors such as working memory capacity or the amount of experience with the stimulus, which is known to change within an individual over time (Anstis and Kim 2011). We also can’t rule out that the duration of the *global* percept may be better reflected in aspects of brain structure rather than function (Kanai et al. 2010, 2011).

### Conclusion

Taken together, our findings show that a constant exercise in interpreting the visual input leaves a trace in our brain even when we are not engaged in any specific task. They show that the anterior IPS shapes the subjective aspects of perception by interacting with nodes of the core DMN and the extrastriate cortex. This interaction may allow an individual to disengage from the sensory input and shorten the time it takes until they can perceive the illusory interpretation of the stimulus. Overall, our results point to the important role of cooperation between a set of areas that are distributed across the key intrinsic networks in shaping the individual’s subjective, illusory aspects of perception.

## Data and code availability statement

Data and code will be made public on the Open Science Framework (OSF) upon manuscript acceptance.

## Notes

### Conflict of interest

None declared.

## Funding

This work was supported by the BioTechMed-Graz Young Research Group Grant to N.Z.

## Acknowledgments

The authors thank Erik Fink, Magdalena Lhotka, and Thomas Zussner for their support in data acquisition. This work was funded by the BioTechMed-Graz Young Research Group Grant Program.

## Notes

### Competing Interest Statement

The authors have declared no competing interest.

## References

Alais D. 2012. Binocular rivalry: Competition and inhibition in visual perception. Wiley Interdiscip Rev Cogn Sci. 3:87–103.

Alderson-Day B, Diederen K, Fernyhough C, Ford JM, Horga G, Margulies DS, McCarthy-Jones S, Northoff G, Shine JM, Turner J, van de Ven V, Van Lutterveld R, Waters F, Jardri R. 2016. Auditory hallucinations and the Brain’s resting-state networks: findings and methodological observations. Schizophr Bull. 42:1110–1123.

Andrews-Hanna JR, Smallwood J, Spreng RN. 2014. The default network and self-generated thought: Component processes, dynamic control, and clinical relevance. Ann N Y Acad Sci. 1316:29–52.

Anstis S, Kim J. 2011. Local versus global perception of ambiguous motion displays. J Vis. 11:13.

Baker DH, Karapanagiotidis T, Coggan DD, Wailes-Newson K, Smallwood J. 2015. Brain networks underlying bistable perception. Neuroimage. 119:229–234.

Birn RM. 2012. The role of physiological noise in resting-state functional connectivity. Neuroimage. 62:864–870.

Boeykens C, Wagemans J, Moors P. 2019. Does task relevance shape the “shift to global” in ambiguous motion perception? J Vis. 19:1–10.

Brascamp JW, van Ee R, Pestman WR, van der Berg A V. 2005. Distributions of alternation rates in various forms of bistable perception. J Vis. 5:287–298.

Britz J, Landis T, Michel CM. 2009. Right Parietal Brain Activity Precedes Perceptual Alternation of Bistable Stimuli. Cereb Cortex. 19:55–65.

Cao T, Wang L, Sun Z, Engel SA, He S. 2018. The independent and shared mechanisms of intrinsic brain dynamics: Insights from bistable perception. Front Psychol. 9:1–11.

Carter OL, Pettigrew JD. 2003. A common oscillator for perceptual rivalries? Perception. 32:295–305.

Chang C, Glover GH. 2009. Effects of model-based physiological noise correction on default mode network anti-correlations and correlations. Neuroimage. 47:1448–1459.

Cichy RM, Chen Y, Haynes JD. 2011. Encoding the identity and location of objects in human LOC. Neuroimage. 54:2297–2307.

Dale AM, Fischl B, Sereno MI. 1999. Cortical Surface-Based Analysis. Neuroimage. 9:179–194.

Deckers RHR, van Gelderen P, Ries M, Barret O, Duyn JH, Ikonomidou VN, Fukunaga M, Glover GH, de Zwart JA. 2006. An adaptive filter for suppression of cardiac and respiratory noise in MRI time series data. Neuroimage. 33:1072–1081.

Devia C, Concha-Miranda M, Rodríguez E. 2022. Bi-Stable Perception: Self-Coordinating Brain Regions to Make-Up the Mind. Front Neurosci. 15:1–20.

Esteban O, Markiewicz CJ, Blair RW, Moodie CA, Isik AI, Erramuzpe A, Kent JD, Goncalves M, DuPre E, Snyder M, Oya H, Ghosh SS, Wright J, Durnez J, Poldrack RA, Gorgolewski KJ. 2019. fMRIPrep: a robust preprocessing pipeline for functional MRI. Nat Methods. 16:111–116.

Fischl B, Van Der Kouwe A, Destrieux C, Halgren E, Ségonne F, Salat DH, Busa E, Seidman LJ, Goldstein J, Kennedy D, Caviness V, Makris N, Rosen B, Dale AM. 2004. Automatically Parcellating the Human Cerebral Cortex. Cereb Cortex. 14:11–22.

Gallagher RM, Arnold DH. 2014. Interpreting the temporal dynamics of perceptual rivalries. Perception. 43:1239–1248.

Glover GH, Li TQ, Ress D. 2000. Image-based method for retrospective correction of physiological motion effects in fMRI: RETROICOR. Magn Reson Med. 44:162–167.

González-García C, Flounders MW, Chang R, Baria AT, He BJ. 2018. Content-specific activity in frontoparietal and default-mode networks during prior-guided visual perception. Elife. 7:1–25.

Gorgolewski K, Burns CD, Madison C, Clark D, Halchenko YO, Waskom ML, Ghosh SS. 2011. Nipype: A flexible, lightweight and extensible neuroimaging data processing framework in Python. Front Neuroinform. 5.

Grassi PR, Zaretskaya N, Bartels A. 2018. A generic mechanism for perceptual organization in the parietal cortex. J Neurosci. 38:7158–7169.

Greve DN, Fischl B. 2009. Accurate and Robust Brain Image Alignment using Boundary-based Registration. Neuroimage. 48:63–72.

Grill-Spector K, Kourtzi Z, Kanwisher N. 2001. The Lateral Occipital Complex and its Role in Object Recognition. Vision Res. 41:1409–1422.

Hampson M, Driesen NR, Skudlarski P, Gore JC, Constable RT. 2006. Brain Connectivity Related to Working Memory Performance. 26:13338–13343.

Hesselmann G, Kell CA, Eger E, Kleinschmidt A. 2008. Spontaneous local variations in ongoing neural activity bias perceptual decisions. Proc Natl Acad Sci U S A. 105:10984–10989.

Kanai R, Bahrami B, Rees G. 2010. Human parietal cortex structure predicts individual differences in perceptual rivalry. Curr Biol. 20:1626–1630.

Kanai R, Carmel D, Bahrami B, Rees G. 2011. Structural and functional fractionation of right superior parietal cortex in bistable perception. Curr Biol. 21.

Kasper L, Bollmann S, Diaconescu AO, Hutton C, Heinzle J, Iglesias S, Hauser TU, Sebold M, Manjaly ZM, Pruessmann KP, Stephan KE. 2017. The PhysIO Toolbox for Modeling Physiological Noise in fMRI Data. J Neurosci Methods. 276:56–72.

Kleiner M, Brainard D, Pelli D, Ingling A, Murray R, Broussard C. 2007. What’s new in Psychtoolbox-3?

Kleinschmidt A, Sterzer P, Rees G. 2012. Variability of perceptual multistability: From brain state to individual trait. Philos Trans R Soc B Biol Sci. 367:988–1000.

Knapen T, Brascamp J, Pearson J, van Ee R, Blake R. 2011. The role of frontal and parietal brain areas in bistable perception. J Neurosci. 31:10293–10301.

Konishi M, McLaren DG, Engen H, Smallwood J. 2015. Shaped by the past: The default mode network supports cognition that is independent of immediate perceptual input. PLoS One. 10:1–18.

Levelt *W*. 1967. Note on the distribution of dominance times in binocular rivalry. Br J Psychol. 58:143–145.

Lorenceau J, Shiffrar M. 1992. The influence of terminators on motion integration across space. Vision Res. 32:263–273.

Mao Y, Kanai R, Ding C, Bi T, Qiu J. 2020. Temporal variability of brain networks predicts individual differences in bistable perception. Neuropsychologia. 142:107426.

Margulies DS, Ghosh SS, Goulas A, Falkiewicz M, Huntenburg JM, Langs G, Bezgin G, Eickhoff SB, Castellanos FX, Petrides M, Jefferies E, Smallwood J. 2016. Situating the default-mode network along a principal gradient of macroscale cortical organization. Proc Natl Acad Sci U S A. 113:12574–12579.

Martinez A, Ramanathan DS, Foxe JJ, Javitt DC, Hillyard SA. 2007. The role of spatial attention in the selection of real and illusory objects. J Neurosci. 27:7963–7973.

Martínez GAR, Parra HC. 2018. Bistable perception: Neural bases and usefulness in psychological research. Int J Psychol Res. 11:63–76.

Moreno-Bote R, Rinzel J, Rubin N. 2007. Noise-induced alternations in an attractor network model of perceptual bistability. J Neurophysiol. 98:1125–1139.

Moreno-Bote R, Shpiro A, Rinzel J, Rubin N. 2010. Alternation rate in perceptual bistability is maximal at and symmetric around equi-dominance. J Vis. 10:1–18.

Murphy C, Jefferies E, Rueschemeyer SA, Sormaz M, Wang H ting, Margulies DS, Smallwood J. 2018. Distant from input: Evidence of regions within the default mode network supporting perceptually-decoupled and conceptually-guided cognition. Neuroimage. 171:393–401.

Murphy K, Birn RM, Bandettini PA. 2013. Resting-state FMRI confounds and cleanup. Neuroimage. 349–359.

Norcia A. 2006. The Best Illusion of the Year Contest.

Pearson J, Westbrook F. 2015. Phantom perception: Voluntary and involuntary nonretinal vision. Trends Cogn Sci. 19:278–284.

Reineberg AE, Andrews-Hanna JR, Depue BE, Friedman NP, Banich MT. 2015. Resting-state Networks Predict Individual Differences in Common and Specific Aspects of Executive Function. Neuroimage. 1:69–78.

Rubin E. 1915. Synsoplevede Figurer. Gyldendalske Boghandel, Copenhagen.

Schaefer A, Kong R, Gordon EM, Laumann TO, Zuo X-N, Holmes AJ, Eickhoff SB, Yeo BTT. 2018. Local-Global Parcellation of the Human Cerebral Cortex from Intrinsic Functional Connectivity MRI. Cereb Cortex. 28:3095–3114.

Schooler JW, Smallwood J, Christoff K, Handy TC, Reichle ED, Sayette MA. 2011. Meta-awareness, perceptual decoupling and the wandering mind. Trends Cogn Sci. 15:319–326.

Seeley WW, Menon V, Schatzberg AF, Keller J, Glover GH, Kenna H, Reiss AL, Greicius MD. 2007. Dissociable intrinsic connectivity networks for salience processing and executive control. J Neurosci. 27:2349–2356.

Shpiro A, Curtu R, Rinzel J, Rubin N. 2007. Dynamical Characteristics Common to Neuronal Competition Models. J Neurophysiol. 97:462–473.

Smallwood J, Bernhardt BC, Leech R, Bzdok D, Jefferies E, Margulies DS. 2021. The default mode network in cognition: a topographical perspective. Nat Rev Neurosci. 22:503–513.

Smallwood J, Tipper C, Brown K, Baird B, Engen H, Michaels JR, Grafton S, Schooler JW. 2013. Escaping the here and now: Evidence for a role of the default mode network in perceptually decoupled thought. Neuroimage. 69:120–125.

Sormaz M, Murphy C, Wang H, Hymers M, Karapanagiotidis T, Poerio G. 2018. Default mode network can support the level of detail in experience during active task states. Proc Natl Acad Sci. 115:9318–9323.

Sterzer P, Kleinschmidt A. 2010. Anterior insula activations in perceptual paradigms: often observed but barely understood. Brain Struct Funct. 214:611–622.

Sterzer P, Kleinschmidt A, Rees G. 2009. The neural bases of multistable perception. Trends Cogn Sci. 13:310–318.

Wallach H, O’Connell DN. 1953. The kinetic depth effect, Journal of Experimental Psychology.

Wang L, Mruczek REB, Arcaro MJ, Kastner S. 2015. Probabilistic maps of visual topography in human cortex. Cereb Cortex. 25:3911–3931.

Wilding M, Körner C, Ischebeck A, Zaretskaya N. 2022. Increased insula activity precedes the formation of subjective Gestalt. Neuroimage. 257:1–30.

Yeshurun Y, Nguyen M, Hasson U. 2021. The default mode network: where the idiosyncratic self meets the shared social world. Nat Rev Neurosci. 22:181–192.

Yeshurun Y, Swanson S, Simony E, Chen J, Lazaridi C, Honey CJ, Hasson U. 2017. Same Story, Different Story: The Neural Representation of Interpretive Frameworks. Psychol Sci. 28:307–319.

Zaretskaya N, Anstis S, Bartels A. 2013. Parietal cortex mediates conscious perception of illusory gestalt. J Neurosci. 33:523–531.

