## Supplementary Material for "Resting-state functional connectivity predictors of subjective visual Gestalt experience"

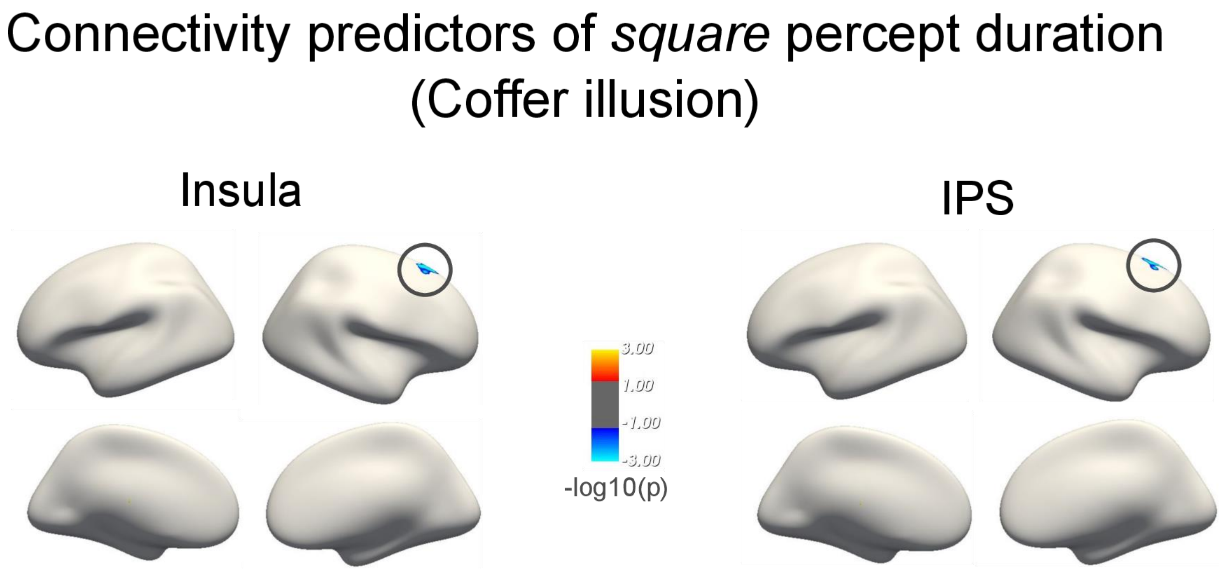


*Figure S1*. Resting-state functional connectivity predictors of the *square* percept duration of the Coffer illusion with both seed regions respectively; p< .01, corrected.

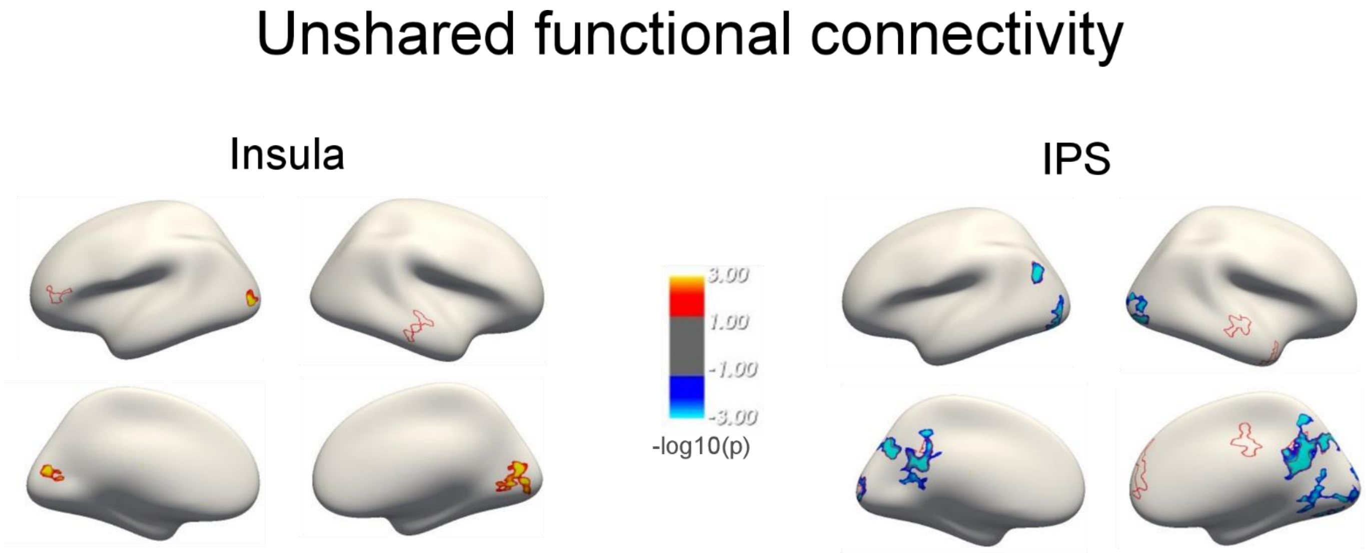


*Figure S2*. Unshared FC for the insula and IPS seed (both seeds were added into the same first-level GLM). Red clusters indicate positive, blue clusters indicate the inverse relationship between connectivity strength and the *local* percept duration. Red outlines indicate seed FC, using a different GLM per seed; both $p<0.01$, corrected.
